# Diffusion MRI Changes in the Healthy Aging Canine Brain

**DOI:** 10.1101/2020.10.05.327205

**Authors:** Erica F. Barry, John P. Loftus, Wen-Ming Luh, Mony J. de Leon, Sumit N. Niogi, Philippa J. Johnson

**Affiliations:** Cornell College of Veterinary Medicine, Cornell University, Ithaca, NY; National Institute on Aging, Baltimore, Maryland; Department of Radiology, Weill Cornell Medicine, New York, NY

**Keywords:** Dog, White matter, DWI, TBSS, FA, AxD

## Abstract

White matter dysfunction and degeneration have been a topic of great interest in healthy and pathological aging. While ex vivo studies have investigated age-related changes in canines, little in vivo canine aging research exists. Quantitative diffusion MRI such as diffusion tensor imaging (DTI) has demonstrated aging and neurodegenerative white matter changes in humans. However, this method has not been applied and adapted in vivo to canine populations. This study aimed to test the hypothesis that white matter diffusion changes frequently reported in human aging are also found in aged canines. The study used Tract Based Spatial Statistics (TBSS) and a region of interest (ROI) approach to investigate age related changes in fractional anisotropy (FA), mean diffusivity (MD), axial diffusivity (AxD) and radial diffusivity (RD). The results show that, compared to younger animals, aged canines have significant decreases in FA in parietal and temporal regions as well as the corpus callosum and fornix. Additionally, AxD decreases were observed in parietal, frontal and midbrain regions. Similarly, an age-related increase in RD was observed in the right parietal lobe while MD decreases were found in the midbrain. These findings suggest that canine samples offer a model for healthy human aging as they exhibit similar white matter diffusion tensor changes with age.

## 1. Introduction

Canine populations are increasingly studied as a translational model for the neuro-mechanisms behind human aging and cognitive decline. Aging canine subjects exhibit similar neuropathology to aging humans, such as the development of beta-amyloid plaques (Aβ) and tauopathy (Cummings et al., 1996, 1993; Head, 2013a; Mazzatenta et al., 2017). Additionally, as in human cohorts a high percentage of dogs exhibit age-related cognitive decline in memory, mood and processing speed related cognitive domains (Cotman and Head, 2008; Head, 2013b). An estimated 60% of older dogs are affected by canine cognitive dysfunction (CCD) (Fast et al., 2013) which is defined by a decline in cognitive function accompanied by cerebral degeneration and decreases in cortical volume (De Angelis et al., 2015; Kimotsuki et al., 2005), similar to that observed in mild cognitive impairment as related to early Alzheimer’s disease in humans (Reisberg et al., 1982; Schütt et al., 2018).

Volumetric MRI studies utilizing voxel based morphometry have reported comparable volumetric changes in canines to those observed in the aging human such as reductions in the hippocampus and frontal, parietal, temporal and occipital lobes (Kimotsuki et al., 2005; Pugliese et al., 2010; Su et al., 1998). The aging canine brain exhibits age-dependent myelin loss (Chambers et al. 2012) similar to the white matter abnormalities associated with cognitive decline in humans. However, to date, white matter microstructure in the aging canine has not been evaluated in vivo.

Diffusion tensor magnetic resonance imaging (DTI), a form of MRI that can quantify restriction of Brownian motion of water molecules in brain tissue, is particularly well suited to study underlying microstructural properties of white matter (Alexander et al., 2007). The tensor based diffusivity metric fractional anisotropy (FA) quantifies anisotropy of diffusion (Basser et al., 1994; Basser and Jones, 2002). In white matter, FA is relatively high due to the underlying axonal structure constraining diffusion orthogonal to the direction of the tract. While diffusion tensor metrics in white matter are influenced by several underlying cellular properties, including axonal density, fiber orientation, and myelination, FA is often interpreted as a general measure of axonal coherence and microstructural integrity (Madden et al., 2009a; Niogi et al., 2007). Other complementary tensor metrics, including mean diffusivity (MD), radial diffusivity (RD), and axial diffusivity (AxD) are commonly utilized to provide additional characterization of molecular diffusion within a voxel. Measures of AxD and RD have been respectively associated with axonal loss and myelination in preclinical models and thus may aid in the interpretation of DW-MRI results (Bennett et al., 2011; Budde et al., 2007; Nair et al., 2005; Song et al., 2002, 2003).

In the human brain, DTI has proven to be an essential tool for detecting and monitoring white matter degeneration in aging and has even shown potential to accurately predict conversion to age-related cognitive impairment (Kruggel et al., 2017). Typically, studies of healthy aging humans report widespread FA decreases and MD increases in white matter that appear to follow an anterior to posterior gradient of neurodegeneration over time (Coutu et al., 2014; Pfefferbaum et al., 2000). Longitudinal studies have found an inverse parabolic relationship between myelination, FA, and aging across the healthy lifespan, which is particularly prominent in the frontal lobe and corpus callosum (Bartzokis et al., 2012; Lebel et al., 2019; Lebel and Deoni, 2018). These findings correspond well with ex vivo reports that indicate age-dependent demyelination (Chambers et al., 2012).

Despite the strengths of DTI in evaluating white matter degeneration in aging and the unique benefits offered by canines as a model of human aging, to our knowledge, no DTI study has reported on in-vivo white matter changes in the aging dog brain. Accordingly, this study aimed to assess white matter changes associated with aging in a canine population using DTI to quantify white matter changes in vivo. Using well-validated tract-based spatial statistic (TBSS) and region of interest (ROI) techniques, we investigated differences in FA, MD, AxD, and RD between neurologically typical younger and old aged canine cohorts. We hypothesize that, consistent with human aging literature (Madden et al., 2009a; Sexton et al., 2011), i) there will be widespread FA and AxD decreases throughout the brain in the aged subjects compared to young controls and associated widespread MD and RD increases and ii) aged associated FA and AxD decreases and MD and RD increases will follow an anterior-posterior pattern with significant differences being more prominent in rostral ROIs.

## 2. Materials and Methods

### 2.1 Subjects

Twenty canine subjects were recruited from research populations (Cornell University College of Veterinary Medicine). Subjects were included if they were evaluated as healthy on physical and neurological examinations. Brachycephalic dogs were excluded in order to limit variation of brain structure between subjects secondary to cranial conformation. The recruited population was divided into two groups according to age: dogs ≥ greater than 10 years and dogs ≤ 7years of age. The aged cohort included 10 subjects (Female = 6, Male = 4) with a median age of 10 (range=10-11 years) years and a median weight of 22 kg (range=20-31kg). The control cohort included 10 subjects (Female = 7, Male = 3) with a median age of 5 (range 4-6 years) years old and a median weight of 13 kg (range=10-30kg). All subjects conformed to mesaticephalic cranial conformation (Carreira and Ferreira, 2015; Hecht et al., 2019; Pilegaard et al., 2017). All aspects of this study were approved by the Cornell University Institutional Animal Care and Use Committee (IACUC protocol number: 2015-0115) (Table 1).

**Table 1.** Results of t-tests conducted to identify significant differences between younger and aged groups for each ROI to explore regional patterns of FA, AxD, MD, and RD changes with age. The single symbol indicates a decrease (-) or increase (+) at the p<0.05 significance level while the double symbol indicates a decrease (--) or increase (++) at the p<0.01 significance level (young=10, old=10).

| Region of Interest |  | Diffusivity Parameter |  |  |  |
| --- | --- | --- | --- | --- | --- |
|  |  | FA | AxD | RD | MD |
| Frontal | Left |  | -- |  |  |
|  | Right |  | - |  |  |
| Sensory-motor | Left |  |  |  |  |
|  | Right |  |  |  |  |
| Cingulate | Left |  |  |  |  |
|  | Right |  |  |  |  |
| Parietal | Left | -- | - |  |  |
|  | Right | -- | - | + |  |
| Temporal | Left |  |  |  |  |
|  | Right | - |  |  |  |
| Occipital | Left |  |  |  |  |
|  | Right |  |  |  |  |
| Corpus Callosum | Genu | - |  |  |  |
|  | Body | -- |  |  |  |
|  | Splenium | -- |  |  |  |
| Other | Fornix | - |  |  |  |
|  | Mid Brain |  | - |  | - |
|  | Cerebellum |  |  |  |  |

### 2.2 MRI Examination

Dogs underwent magnetic resonance imaging (MRI) under general anesthesia performed by a board-certified veterinary anesthesiologist. Dogs were premedicated with Dexmedetomidine (3 mcg/kg) and Dexdomitor (0.5 mg/ml, Zoetis Inc, Kalamazoo, MI), and induced to general anesthesia with propofol (3.2-5.4 mg/kg Sagent Pharmaceuticals, Schaumburg, III), and then intubated. They were maintained under anesthesia with inhalant isoflurane and oxygen with a dexmedetomidine continuous rate infusion (1 mcg/kg/hr Dexdomitor 0.5 mg/ml, Zoetis Inc, Kalamazoo, MI). MRI was performed in a 3.0T GE Discovery MR750 (GE Healthcare, Milwaukee, WI) whole-body scanner (60 cm bore diameter), operating at 50 mT/m amplitude and 200 T/m/s slew-rate. Subjects were placed in dorsal recumbency with their head centered in a 16-channel medium flex radio-frequency coil (NeoCoil, Pewaukee, WI 53072 USA). A high-resolution T1-weighted 3D inversion-recovery fast spoiled gradient echo sequence (Bravo) was performed in each subject with the following parameters; isotropic voxels 0.5mm^3^, TE=3.6ms, TR=8.4ms, TI=450ms, NEX=3, 12° flip angle, and acquisition matrix size=256×256. Diffusion tensor images were acquired in the transverse plane (TR=7000ms, TE=89.6ms, flip angle=90°, isometric voxel size of 1.5×1.5×1.5mm, in-plane field of view=135×135mm, matrix size 90×90 with 60 gradient directions, b=800 s/mm^2^ and a single unweighted (b=0) diffusion image.

### 2.3 DW-MRI Preprocessing

DWI images were corrected for noise (Veraart et al., 2016), phase distortion (Andersson et al., 2003; Smith et al., 2004), removing Gibbs artifact (Kellner et al., 2016), eddy current distortion and motion correction (Andersson and Sotiropoulos, 2016) using the FSL (https://fsl.fmrib.ox.ac.uk/) and MRTrix (https://www.mrtrix.org) software packages. Diffusion tensors were then modeled using FSL’s *dtifit* from the FSL diffusion toolbox (Behrens et al., 2007, 2003). Each tensor was defined by three principal eigenvalues (i.e.,. λ_1_, λ_2_, λ_3_). Tensor maps were then calculated for fractional anisotropy (FA; 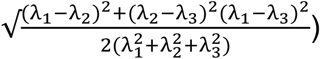, mean diffusivity (MD; (λ_1_ + λ_2_ + λ_3_)/3), radial diffusivity (RD; λ_2_ + λ_3_)/2), and axial diffusivity (AxD; λ1) (Basser and Jones, 2002; Beaulieu, 2002). Diffusion tensor maps for each diffusivity parameter were generated for each subject and visually inspected to ensure the quality of between volume registration, orientation, and preprocessing.

### 2.4 TBSS Analysis

A modified tract-based spatial statistic (TBSS) analysis was conducted (Jbabdi et al., 2010) Smith 2006?. Subjects’ FA images were first organized according to TBSS protocols (https://fsl.fmrib.ox.ac.uk/fsl/fslwiki/TBSS/UserGuide). The first step of TBSS processing was conducted according to human protocols using *tbss_1_preproc*, to slightly erode each FA image and set the end slices of each FA image to zero to eliminate outliers from tensor fitting. *tbss_2_reg*, was then used to perform nonlinear registration from each subject to every other subject in order to find the most representative subject of the group. Once all the combinations of registrations were complete, and the target subject was chosen using *tbss_3_postreg*.

From this step forward modifications from the human pipeline were implemented. Since there is no standardized template available for canine populations at this time, Advanced Normalization Tools (ANTs) nonlinear registration symmetric image normalization (SyN) (Avants et al., 2008) was used to register each subject’s FA image to the target FA image. The target space FA images were then concatenated into a single 4D file that was then averaged using *fslmaths* to create a mean FA image. This mean FA image was then masked at a lower threshold of 0.2 and an upper threshold of 0.8 to create a mean FA skeleton which was then binerized to create a FA skeleton mask to isolate voxels in subsequent processing. The FA skeleton mask was then applied to each subject’s FA image in target space. The resulting subject’s FA skeletons were concatenated into a single 4D FA skeleton image. The other tensor measures, MD, RD, and AxD, were processed according to the same steps outlined above and extracted using the FA skeleton. FA, MD, RD and AxD values at the location of the FA skeleton mask were then exported for statistical analysis.

### 2.5 Statistics

Permutation testing using FSL’s *randomise* tool was used to conduct independent t-test to evaluated differences in the groups using both Threshold Free Cluster Enhancement (TFCE) and Family-Wise Error correction (FWE) to control for multiple comparisons (Salimi-Khorshidi et al., 2011; Smith and Nichols, 2009; Winkler et al., 2014). A two-group design with a continuous covariate interaction test adjusted for the effect of weight (an estimator of breed) on the design matrix (Barnett et al., 1975). A two-group design with continuous covariate interaction test was implemented to adjust for the effect of weight (an estimator of breed) on the design matrix (Barnett et al., 1975). Due to nonparametric nature of the data, Mann-Whitney U tests were used to test differences in FA, MD, AxD, and RD between the young and aged groups. Mann-Whitney U tests were used across 20 regions of interest within the white matter skeleton to investigate regional patterns of aging (Table 1.). False discovery rate (FDR) was applied to correct for multiple comparisons (Benjamini and Hochberg, 1995).

### 2.6 ROI Analysis

To assess the potential regional pattern of diffusivity changes, regions of interest (ROI) were selected to be tested across age groups. A stereotaxic canine cortical atlas was used to apply specific ROIs for each hemisphere that included the cingulate, frontal, occipital, temporal, sensory-motor and parietal (Johnson et al., 2020). The same atlas was used to source the following structures: the fornix, midbrain, cerebellum, genu, body, and splenium. These regions were selected to represent human brain regions most effected by age and test for an anterior-posterior effect. The masks from the atlas were linearly registered to the target space from the TBSS analysis and manually inspected by two investigators for quality assurance. The ROI masks were applied to the mean FA skeleton masked subjects’ tensor maps in target space to ensure sampling from white matter voxels only. The delineation of these regions is visualized in Figure 1. These masks were then used as ROIs across subjects in target space in order to find the mean for each diffusion tensor measure (FA, MD, AxD, and RD) across subjects using *fslstats* (Jenkinson et al., 2012).

**Figure 1.**
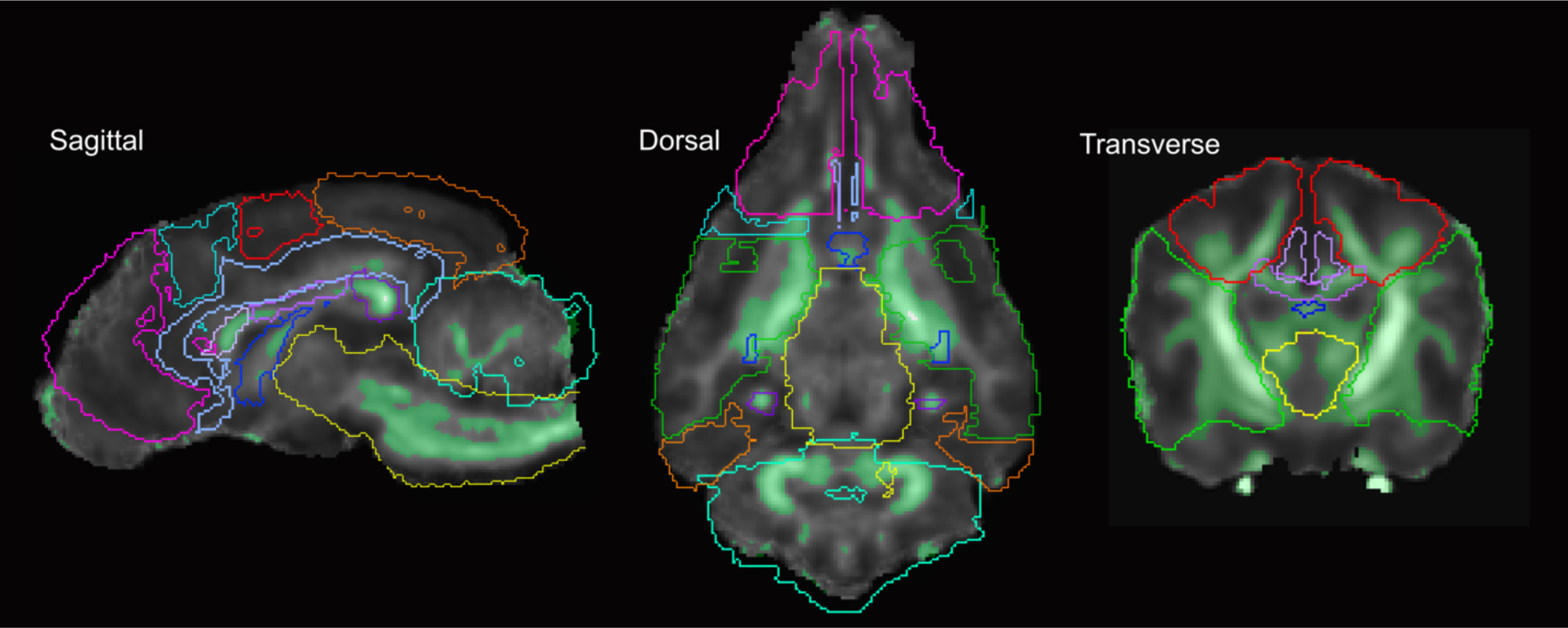
Sagittal, axial and transverse views of the delineation of lobes for ROI analysis. The frontal (magenta), sensory-motor (light blue), cingulate (lavender), parietal (red), genu (light purple), body of corpus callosum (purple), splenium (dark purple), fornix (dark blue), midbrain (yellow), occipital (orange), cerebellum (teal) and temporal (green) ROI mask outlines are overlaid on a template FA and white matter skeleton mask (light green).

## 3. Results

### 3.1 Voxel-based analysis with TBSS

Compared with control canine subjects, the TBSS analysis showed widespread significant decreases in FA particularly in the medial aspects of the white matter skeleton in aged subjects, Figure 2. Furthermore, compared with control canine subjects, aged canines showed significant decreases in AxD in several white matter regions Figure 2. No significant differences in the white matter RD or MD between control and aged subjects were observed.

**Figure 2.**
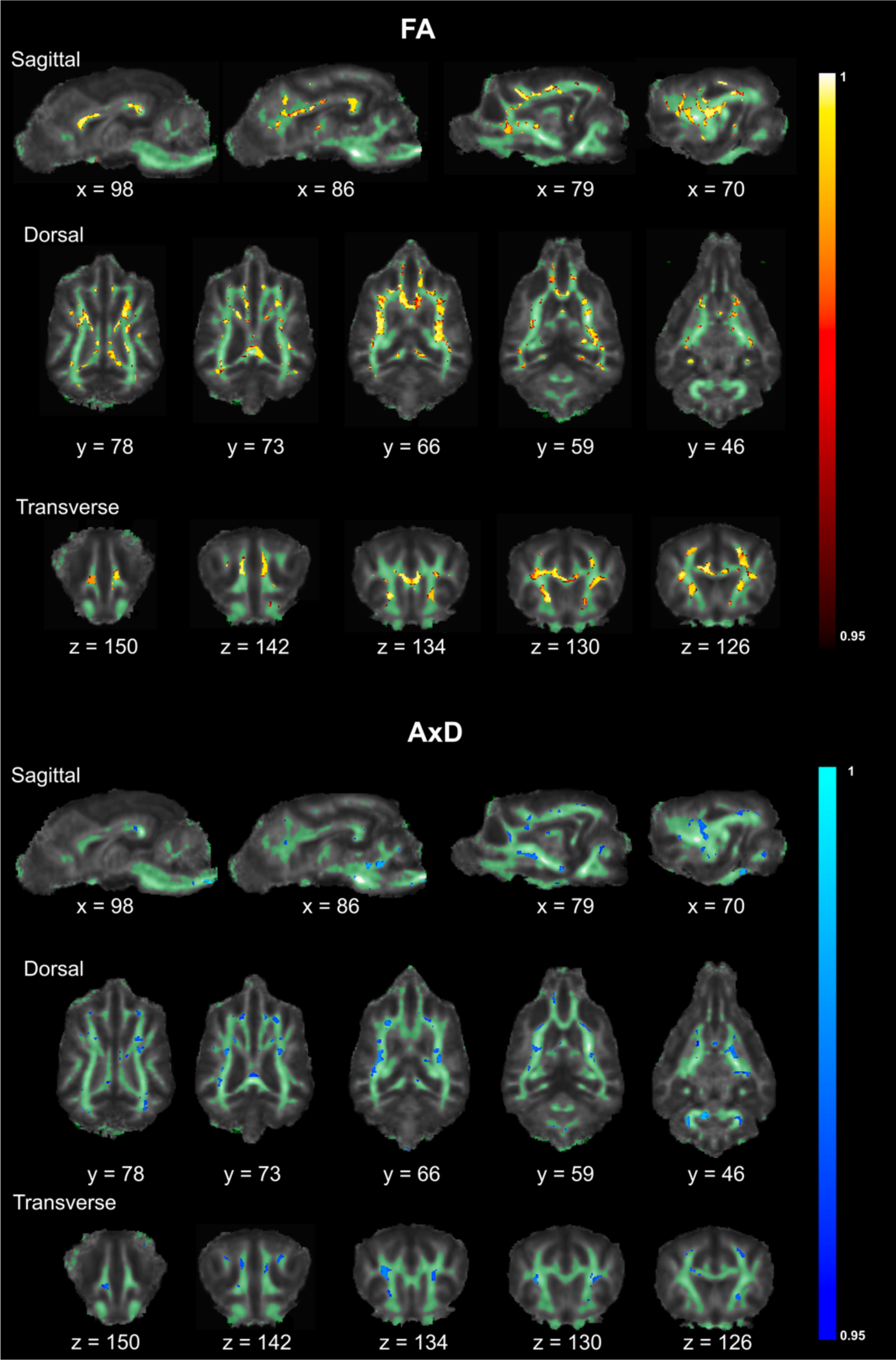
Light green overlay of the average white matter mask over sample FA template showing FA (heatmap) and AxD (cool-map) levels of significant decreases from p<0.05(0.95) to p<0.0001 (1) featuring slices in sagittal, axial and transverse plans to show TBSS analysis results.

### 3.2 ROI analysis

An ROI analysis of 20 regions was conducted for all tensor measures masked within the white matter skeleton. Boxplots showing significant differences in diffusivity parameters in bilateral and unilateral ROIs between control and aged subjects are shown in Figure 3. A Mann-Whitney U-test showed significant decreases in FA in the aged group compared with the young group in the left parietal (*Mann Whitney* U=6, old median=0.30, young median=0.34, p<0.05), right parietal (*Mann Whitney* U=3, old median=0.30, young median=0.36, p<0.001), and right temporal regions (*Mann Whitney* U=21, old median=0.34, young median=0.37, p<0.05). Bilateral significant decreases in the aged group were observed in the genu (*Mann Whitney* U=16, old median=0.34, young median=0.39, p<0.05), body (*Mann Whitney* U=8, old median=0.26, young median=0.30, p<0.05) and splenium of the corpus callosum (*Mann Whitney* U=11, old median=0.32, young median=0.40, p<0.05) as well as the fornix (*Mann Whitney* U=21, old median=0.34, young median=0.37, p<0.05). Significant decreases in AxD were also observed in the aged group compared with the young group in the left frontal (*Mann Whitney* U=12, old median=0.00102, young median=0.00107, p<0.05), right frontal (*Mann Whitney* U=23, old median=0.00101, young median=0.00105, p<0.05), left parietal (*Mann Whitney* U=19, old median=0.00103, young median=0.00106, p<0.05), right parietal (*Mann Whitney* U=21, old median=0.00102, young median=0.00106, p<0.05), and bilateral midbrain (*Mann Whitney* U=20, old median=0.00131, young median=0.00137, p<0.05) regions. Significance levels are reported in Table 1.

**Figure 3.**
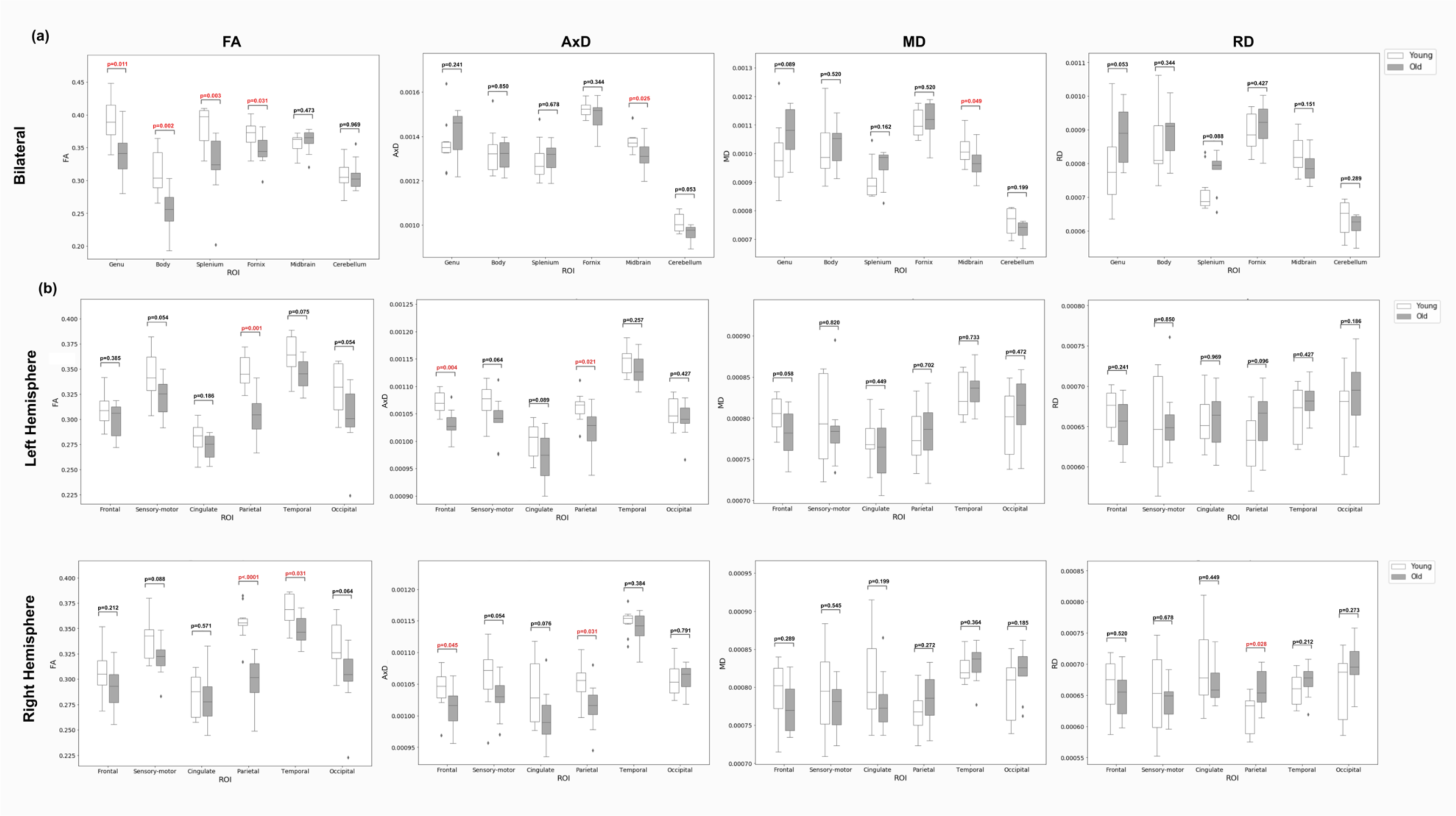
Boxplots featuring bilateral FA, AxD, MD and RD age differences with significance levels highlighted in red featuring the relevant p-values (a). Boxplots featuring unilateral FA and AxD age differences with significance levels highlighted in red featuring the relevant p-values (b). Darker boxes indicate old subjects and white boxes indicating younger subjects. ROIs are ordered in an anterior to posterior fashion and columns are arranged by diffusion tensor measures starting with FA and continuing with AxD, MD and RD.

A significant increase in RD in the young group (N=10) compared with the old group (N=10) in the right parietal region (Mann Whitney U=79.5, old median=0.00079, young median=0.00077, p<0.05) was observed, while a significant decrease in MD was observed in the midbrain of the older group (Mann Whitney U=24, old median=0.00097, young median=0.0010, p<0.05).

## 4. Discussion

This study aimed to investigate if canine populations show age-associated changes in diffusion tensor metrics similar to those previously observed in humans. Using TBSS, we observed significant decreases in FA and AxD in aged canine subjects compared to younger canines. Secondary ROI analysis found widespread decreases in white matter FA and AxD associated with age bilaterally in the parietal regions with a corresponding increase in RD in the right parietal region. Other tensor metric decreases with age appeared to be metric specific such as the FA decrease in the corpus callosum, fornix and right temporal region and AxD decrease in the frontal and midbrain regions. These specific tensor metric results suggest a driving axonal degeneration or loss which may differentiate normal aging from typical demyelination findings with pathological aging.

The widespread age related FA decreases found in the TBSS analysis is consistent with the majority of human aging studies (Abe et al., 2008; Borowski et al., 2018; De Groot et al., 2016; Sullivan and Pfefferbaum, 2006). Corresponding AxD decreases found in the current analysis indicate AxD as the major component driving FA changes in these overlapping regions. The decrease in AxD but the lack of RD increase in these regions may be an indication of histological changes associated with AxD such as neuronal loss (Song et al., 2003). These other results suggest AxD measures may serve as an imaging biomarker to distinguish normal aging from pathological demyelination, which has been associated with RD increases (Gatto et al., 2018; Madden et al., 2012, 2009b; Nir et al., 2013). Conversely, increased white matter MD with age has been observed in many human aging studies (Kantarci et al., 2017; Lebel et al., 2012), but not in the current study. Since MD is determined by the mean of the 3 eigenvalues the observed decrease in AxD and a small increase in the other 2 eigenvalues (constituting RD) could offset an expected MD decrease. Another explanation for this is that because MD because a general measure of diffusivity, this measure may have been affected by a variety of underlying aspects of the neuronal environment rather than a specific neuromechanism. Therefore there is potential for multiple underlying neuromechanisms in which MD is sensitive too could have opposing effects on the MD measure and reduce the overall measure

Though FA decreases were found to be widespread in the TBSS analysis, the ROI analysis located decreases in the parietal lobes, right temporal, fornix and the corpus callosum. These results are consistent with human aging studies which found significant decreases in the corpus callosum specifically (Abe et al., 2002; Bennett et al., 2017; Frederiksen, 2013; Sullivan and Pfefferbaum, 2006). In regards to ROI results of this study, aging networks including the fronto-parietal, and default mode network could also be implicated due the specific ROIs FA decreases in our study being found in comparable regions within those networks including the frontal, and parietal regions (Betzel et al., 2014; Brown, 2017). Similarly, our results are consistent with canine aging studies finding cortical atrophy with age and beta-amyloid plaque accumulation associated with behavioral changes including drinking, appetite, social orientation, day-night rhythm and memory in the parietal regions (Rofina et al., 2006; Head et al., 2011). The significant decreases in the parietal lobe is also consistent with findings from human aging literature which have found overactivation in the parietal lobe on a series of attention tasks (Madden 2007; Davis 2008) as well as reduced volume, metabolic decline and cortical thinning (Greenwood et al., 2000). The fronto-parietal network hub disruption is suggested to be a potential mechanism for these changes though these findings need further replication in the human literature (Greenwood et al., 2000). Future studies in canine subjects could seek to use atlases to further delineate the parietal and frontal lobes into sections for hypothesis driven investigations of this network.

Though significant decreases in AxD and FA were found in the frontal and parietal lobes our results did not show a clear anterior-posterior gradient associated with aging. The cross-sectional nature of this study potentially was not sensitive enough to capture diffusion gradient changes that have been shown to vary across lifespan (Coutu et al., 2014; Pfefferbaum et al., 2000). Future studies could include larger samples and more groups of a similar breed of various ages to compare across the lifespan or additionally could conduct longitudinal studies following a reasonably sized cohort across the canine lifespan with multiple timepoints per year to account for the rapid aging of canine populations compare to humans. Additionally, this study faces limitations related to the interpretation of diffusion tensor measures. This is due to the fact that numerous microstructural elements in addition to fiber orientation within a voxel (crossing, fanning, or kissing fibers) may effect these measures (Wheeler-Kingshott and Cercignani, 2009); (Tournier et al., 2012). Despite these limitations, diffusion measures of FA, MD, AxD, and RD remain the most commonly reported metrics in human literature. Because no studies have previously examined these measures in vivo within the canine population, we opted to explore the most commonly reported metrics for comparison with the majority of human aging diffusion literature. However, new acquisition techniques such as Diffusion Kurtosis Imaging (Jensen et al., 2005) or more complex representations of the 3D diffusion profile such as HARDI (Alexander et al., 2002) and Diffusion Spectrum Imaging (Wedeen et al., 2005) are being applied to human samples in order to to resolve intra-voxel tract orientations. Similarly, advanced post-processing methods such as BEDPOSTX (Woolrich et al., 2008) and constrained spherical deconvolution (Johansen-Berg and Behrens, 2014; Tournier et al., 2019) can be used to compute multiple fiber orientation distributions unlike DTI which is limited to modeling single fiber orientation. Combining diffusion measurements with biophysical profile models such as NODDI (Zhang et al., 2012) can potentially increase specificity of our interpretation through additional metrics such as neurite density and orientation dispersion. Taken together, the age-related changes in DTI measures in this canine sample indicate the canine as a valuable translational model of human aging.

In conclusion, this study aimed to investigate difference in diffusion tensor metrics between aged and younger control canine subjects. Decreases in measures of FA in the aged canine group corresponded well to consistent findings in the human literature. Although more research is needed to investigate aging trajectories for in vivo canine samples and the effects of aging on specific networks of interest, the canine population has been shown to have excellent potential as a model for human aging.

## Acknowledgments

We would like to acknowledge the assistance of Zonia Clancy, Carol Frederick and Nora Mathews for their assistance in handling and anesthesia during magnetic resonance imaging. We would also thank the Vaika foundation and the Bowman, Boesch and Cheetham Labs based at Cornell University College of Veterinary Medicine for contributing subjects for this study.

## Disclosure Statement

There were no actual or potential conflicts of interest in regard to this study.

## Notes

### Competing Interest Statement

The authors have declared no competing interest.

